# Lysosome-Related Organelles Orchestrate Guanine Crystal Formation in Pigment Cells

**DOI:** 10.64898/2026.01.05.697611

**Authors:** Anna Gorelick-Ashkenazi, Yuval Barzilay, Tali Lerer-Goldshtein, Tsviya Olender, Zohar Eyal, May Glaser, Yonatan Broder, Nadav Mishol, Rachael Deis, Merav Kedmi, Dvir Gur

## Abstract

Iridosomes, the guanine crystal-forming organelles of pigment-producing iridophores, are among the most versatile, visually striking yet mechanistically uncharacterized organelles in vertebrate biology. Lysosome-related organelles (LROs) support cell type-specific functions by adapting endolysosomal pathways for specialized roles. Here, we show that iridosomes represent a previously unrecognized subtype of LROs. Using transcriptomic profiling of zebrafish iridophores, CRISPR-Cas9-mediated gene disruption, and cryogenic transmission electron microscopy, we define the molecular program underlying iridosome biogenesis. Iridosomes have evolved unique adaptations for crystal growth while retaining core features of other LROs. Key regulators, including RAB32a, AP3M2, and HPS5, are essential for crystal formation, with gene knockouts causing reduced crystal number, altered morphology, and distinct maturation defects. We further identify hallmark LRO features in iridosomes, including intraluminal vesicles and pH-regulated developmental transitions. Cross-species transcriptomic analysis confirms that iridosomes share an LRO signature across vertebrates, including teleost fish and reptiles, suggesting ancient evolutionary origins. These findings establish iridosomes as crystalline LROs and as a model for investigating how cells construct structurally specialized organelles through coordinated trafficking, acid-base regulation, and crystallization, with implications for LRO evolution and human disease.

## Introduction

Iridosomes, the guanine crystal-forming organelles of neural crest–derived iridophores^1,2^, are among the most visually striking yet poorly characterized organelles in vertebrate biology^3,4^. These membrane-bound compartments generate structural color through tightly packed guanine crystals^2,5–7^, enabling camouflage, signaling, and pattern formation across a wide range of species, including fish, amphibians, reptiles and crustaceans^8–20^. Despite their broad functional relevance, the developmental origin, molecular regulation, and classification of iridosomes remain largely unresolved. In contrast, melanosomes, another class of pigment organelles in neural crest–derived melanophores, are well established as Lysosome-related organelles (LROs) and have served as a foundational model for organelle specialization^21–25^. Iridosomes exhibit an extraordinary degree of control over crystal size, shape, and organization, surpassing current manmade capabilities and represent a compelling model for understanding how cells orchestrate crystal formation within membrane-bound organelles^6,14,26–28^. However, whether iridosomes constitute an unrecognized branch of the LRO family, and what molecular machinery orchestrates their formation, has remained an open question.

LROs are a diverse group of cell type-specific compartments derived from the endosomal-lysosomal system^29,30^. Despite their shared origin, LROs exhibit a remarkable diversity in structures, compositions, and functions reflecting the specialized needs of different cell types^31^. For instance, melanosomes in pigment cells synthesize and store melanin for photoprotection^22,23,32^, platelet dense granules release pro-adhesive factors to support homeostasis^33^, and Weibel-Palade bodies in endothelial cells store and secrete proteins vital for vascular health^34^. Additionally, LROs play crucial roles in immune responses, such as the cytotoxic granules of natural killer cells and cytotoxic T lymphocytes that release effector proteins to target pathogens^35^.

The biogenesis and maintenance of LROs are governed by a complex molecular network involving key regulators such as Rab GTPases, adaptor protein (AP) complexes, biogenesis of lysosome-related organelle complexes (BLOC), and vacuolar protein sorting (VPS) machinery^36^. Among these, RAB32 and RAB38 play pivotal roles in vesicular trafficking and cargo delivery, ensuring the specialized maturation of LROs^37^. The BLOC complexes particularly BLOC-1, BLOC-2, and BLOC-3-interact with Rab GTPases to facilitate the targeted transport of specific cargoes to LROs^22,38,39^.

A common hallmark of many LROs is dynamic pH modulation during maturation. For instance, Weibel-Palade bodies, derived from the Trans-Golgi network, undergo acidification^40^, while endosome-derived melanosomes are initially acidic and experience a pH increase as they mature^30,41^. Similarly, recent findings indicate that iridosomes, organelles responsible for the formation of guanine-based crystals, also undergo a pH shift from acidic to neutral during their maturation^42^.

Defects in LRO formation or function are linked to several genetic disorders, including Hermansky-Pudlak syndrome and Chediak-Higashi syndrome, which manifests in conditions such as albinism, immune dysfunction, and bleeding abnormalities due to defective LRO biogenesis in melanosomes, cytotoxic granules, and platelet granules^21,43^. The versatility of LROs lies in their ability to integrate endosomal and lysosomal components to create functionally specialized microenvironments within the cell^4,29–31^. Better understanding LROs provides insight into how cells rewire standard trafficking pathways to serve highly specific physiological roles.

In this study, we utilized zebrafish (*Danio rerio*) larvae as a model system for iridosome development^18,26,27^, taking advantage of the rapid emergence of hundreds of these crystal-forming organelles in the eyes and skin during early development-a process essential for camouflage and visual function^19,44,45^. Through a combination of transcriptomic profiling of isolated iridophores, genetic perturbations, cryogenic transmission electron microscopy (cryo-TEM), live imaging, and ultrastructural investigations, we dissect the cellular origin and molecular regulation of iridosome biogenesis. Our findings reveal that iridosomes are a specialized type of LRO. Specifically, we demonstrate the involvement of key regulators, including Rab GTPases, adaptor protein complexes, and Hermansky-Pudlak syndrome proteins, which collectively orchestrate the formation and maturation of these optically functional crystals. Beyond defining iridosomes as LROs, our study provides new perspectives on how organelle identity is established within specific cell types, and how large numbers of complex organelles are rapidly assembled and matured during early vertebrate development.

To place our findings in a broader evolutionary framework, we performed a comprehensive metadata analysis of publicly available transcriptomic datasets from multiple species. This analysis revealed that iridosomes are conserved as LROs across a range of taxa, including other teleost fish such as medaka (*Oryzias latipes*)^16^, as well as reptiles like the leopard gecko (*Eublepharis macularius*)^46^. This cross-species conservation suggests that the molecular mechanisms uncovered in zebrafish are likely to represent fundamental principles of iridosome biogenesis, underscoring their evolutionary significance and raising the possibility of conserved roles for iridosomes in visual and pigmentation systems across vertebrates.

## Results

### Transcriptomic profiling of zebrafish iridophores reveals key LRO pathways in iridosome biogenesis

Zebrafish utilize the unique optical properties of guanine crystals for a range of functions. These crystals, distributed throughout their skin and eyes (**Fig. 1A**), play a crucial role in coloration and patterning, contributing to camouflage, acting as light barriers, and enhancing visual sensitivity, particularly in low-light environments^19,44,47–51^. In the developing zebrafish eye, intense crystal formation begins early^42,52^ to support high visual acuity, reaching a peak around 5 days post-fertilization (dpf), when large regions of the iris become densely populated with iridophores filled with light-reflecting crystals (**Fig. 1B–E**). Within these iridophores, each guanine crystal forms inside a membrane-bound organelle (**Fig. 1B**), with numerous crystals typically assembling within hours (**Fig. 1F-J** and Supplementary Movies **S1-S2**).

**Figure 1.**
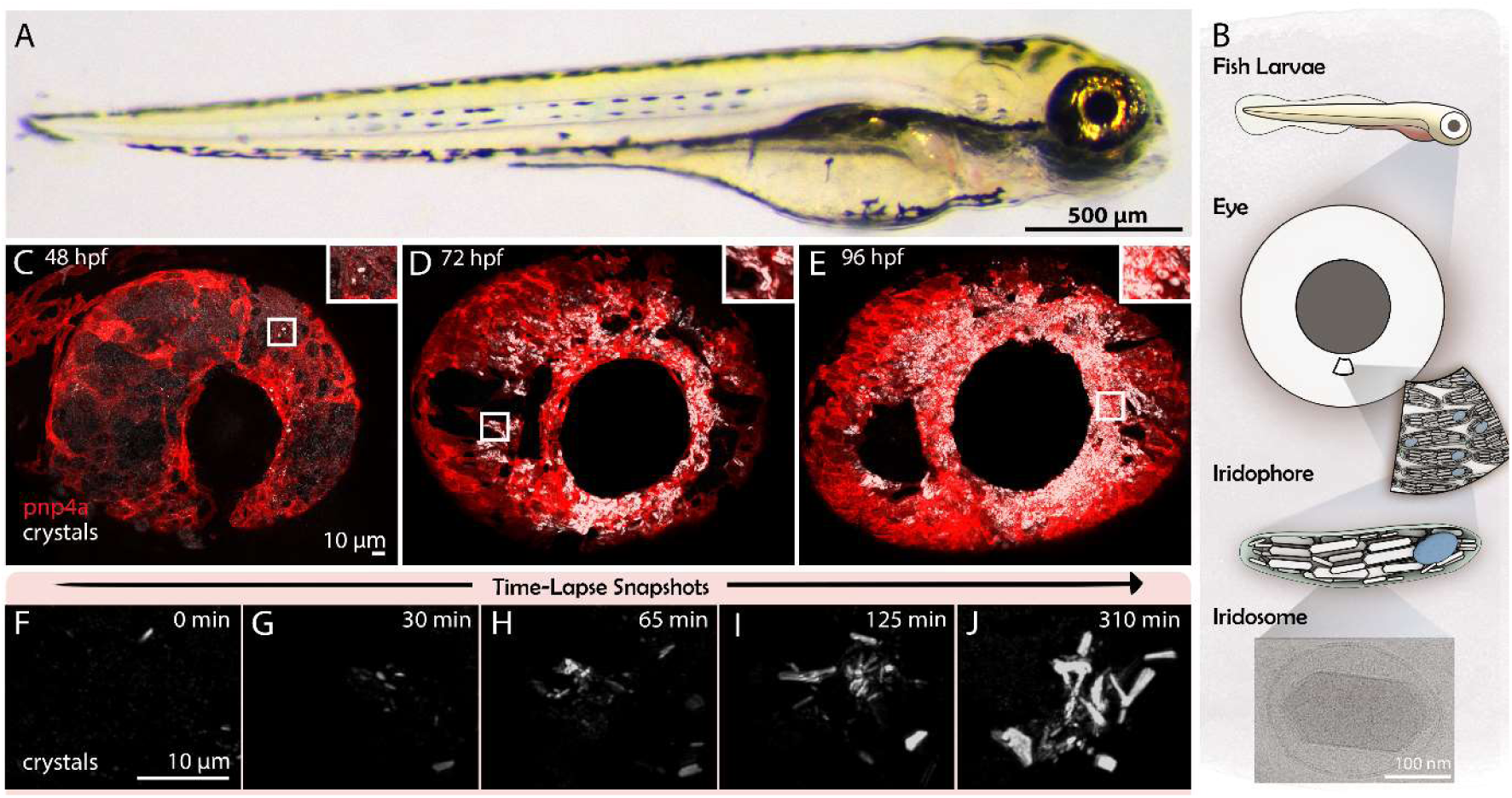
Crystal formation dynamics and development in zebrafish eye iridophores. **(A)** Brightfield image of a 5 dpf zebrafish larva showing the highly reflective iris produced by guanine-based crystals in the eye. (**B**) Schematic of the spatial arrangement of iridophores in the larval eye and the guanine-containing iridosomes within them. Bottom: cryo-electron microscopy image of a single membrane-bound iridosome containing a guanine crystal. (**C–E**) Maximum intensity projections (MIPs) from confocal imaging of *pnp4a*:PALM-mCherry^51^ transgenic larvae at 48, 60, and 72 hpf, revealing the progressive differentiation of iridophores and the accumulation of intracellular crystals during development. Iridophore membranes are labeled in red, while guanine crystal reflectance is shown in white. Insets highlight higher magnification views. (**F–J**) Time-lapse confocal images of a single iridophore in the developing larval eye over 3 hours. Initially, only a few guanine deposits are visible. As development proceeds, crystal number increases rapidly, reflecting active biogenesis of iridosomes during this developmental window.

To investigate the origin and biogenesis of crystal-forming iridosomes, we isolated iridophores from zebrafish larvae at 5 dpf. Fluorescence-activated cell sorting (FACS) followed by transcriptomic analysis was used to compare iridophores to a general, heterogeneous population of cells. We identified 1222 significantly upregulated genes specific to iridophores (**Fig. 2A**), including established iridophore marker genes such as *pnp4a, tfec, slc2a15a, gmps, gpnmb, and alx4a* (**Fig. 2B**)^47,53–57^. Functional enrichment analyses using Metascape^58^ highlighted key pathways enriched in iridophores, particularly the pigment granule organization as well as different lysosome-related pathways (**Fig. 2C, Supplementary Fig. 1C**).

**Figure 2.**
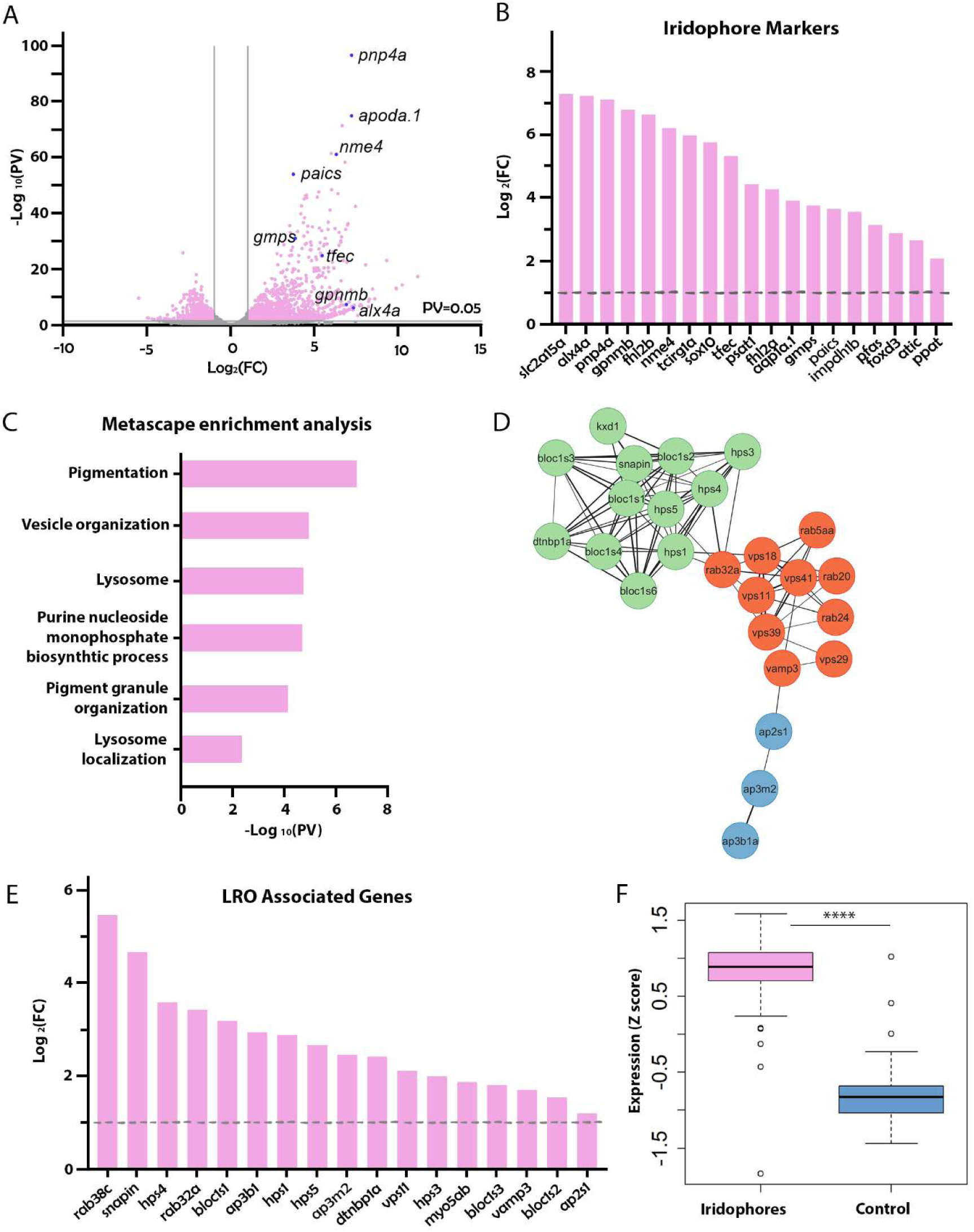
Transcriptomic profiling of zebrafish iridophores reveals lysosome-related organelle signatures. (**A**) Volcano plot showing differentially expressed genes between sorted iridophores (n = 3) and a heterogeneous control cell population (n = 3) from zebrafish larvae at 5 dpf. Genes with P < 0.05 and fold change > 2 are highlighted in pink. Selected iridophore markers are highlighted by blue dots. (**B**) Bar plot illustrating elevated expression of canonical iridophore marker genes, including *pnp4a*, *tfec*, and *gmps*. (**C**) Gene pathways enrichment analysis performed using Metascape^58^, showing significant upregulation of pathways related to lysosomes, vesicle organization, and pigment granule biogenesis (pink bars). (**D**) STRING^63^ (https://string-db.org/) interaction network of LRO-associated genes significantly upregulated in iridophores. The network was clustered using the k-means algorithm into three distinct clusters: BLOC and HPS complexes (green), Rab and VPS proteins (red), and AP complexes (blue) and visualized using Cytoscape^76^. (**E**) Bar graph showing the over expression levels of a consensus set of LRO marker genes in iridophores. **(F)** Distribution of the standardized LRO gene expression in iridophores versus control cells. Iridophores show a significantly higher expression of LRO genes, supporting their classification as LRO-bearing cells. Significance determined by Student’s test, two tailed, unpaired, ****P< 0.0001.

To elucidate the molecular mechanisms underlying iridosome biogenesis, we focused on the machinery and regulatory components implicated in LRO formation. Our transcriptomic analysis revealed an enrichment of over 30 known LRO regulators^22^ that were upregulated in iridophores (**Fig. 2D, E and Supplementary Fig. 1D**). These included members of the Rab GTPase family (such as *rab38*, *rab32*, and *rab5*)^36,37^, adaptor protein complexes (AP) including *ap3* and *ap2* subunits^59,60^, and multiple BLOC complexes components (particularly *bloc1s1* and *bloc1s4*)^36,38^. Additionally, we identified multiple Hermansky-Pudlak syndrome (HPS) proteins (*hps1*, *hps3*, *hps4*, and *hps5*)^7,21,22,36^, as well as various subunits of vacuolar ATPase (V-ATPase)^61^ and vacuolar protein sorting-associated proteins (VPS)^21,36^ (**Fig. 2D and E and Supplementary Fig. 1A**).

To quantify the overall enrichment of LRO genes, we compared the expression of a consensus set of 27 LRO markers^21,22,36,62^ in iridophore samples versus the control samples (**Supplementary Fig. 1B and Supplementary Table. 1)**. Iridophores displayed a significantly elevated expression of LRO genes relative to the control samples (Student’s t-test, two tailed, unpaired, P< 0.0001) (**Fig. 2F, Supplementary Fig. 1B**), strongly suggesting that the iridosomes within iridophores represent a class of LROs.

To understand the functional relationships and connectivity among the upregulated LRO-related genes, we performed STRING analysis^63^. Using the k-means algorithm, we clustered the network into three clusters showing a high degree of connectivity between the genes: one centred around VPS and Rab subunits, a second around BLOC and HPS proteins, and a third includes the AP complexes (**Fig. 2D**). The observed network architecture highlights a coordinated regulatory framework likely governing iridosome biogenesis, closely resembling mechanisms previously described for other LROs, including the closely related melanophores^22^.

In addition to known LRO regulators, our analysis revealed a subset of genes that may represent iridophore-specific adaptations to LRO biology. These include *rab20*, *rab3il1*, and *rabl6b* (**Supplementary Fig. 1A)**, which were selectively upregulated in iridophores but have not been previously associated with melanosome biogenesis. Their expression patterns suggest potential roles in iridosome-specific membrane trafficking, cargo delivery, or organelle–organelle communication. These candidates may reflect functional specialization associated with the unique demands of intracellular guanine crystallization.

### Differential roles of *rab32a* and *ap3m2* in regulating crystal morphology and abundance

Given the notable upregulation of LRO regulators in iridophores, we hypothesized that loss of function of specific LRO-associated genes would significantly impair iridosome biogenesis and guanine crystal formation. To test this, we used clustered regularly interspaced short palindromic repeats (CRISPR)–Cas9 genome editing to generate mutants targeting two key nodes identified in our network analysis (**Fig. 2D**). Specifically, we disrupted *rab32a*, a Rab GTPase known to regulate LRO maturation by coordinating vesicular trafficking and cargo delivery^37,39,64^, and *ap3m2*, the μ3^59,60,65–67^ subunit of the adaptor protein complex AP-3, which mediates cargo sorting from endosomes to LROs^39^. Both mutations resulted in premature stop codons predicted to produce non-functional proteins (**Fig. 3 and Methods**).

**Figure 3.**
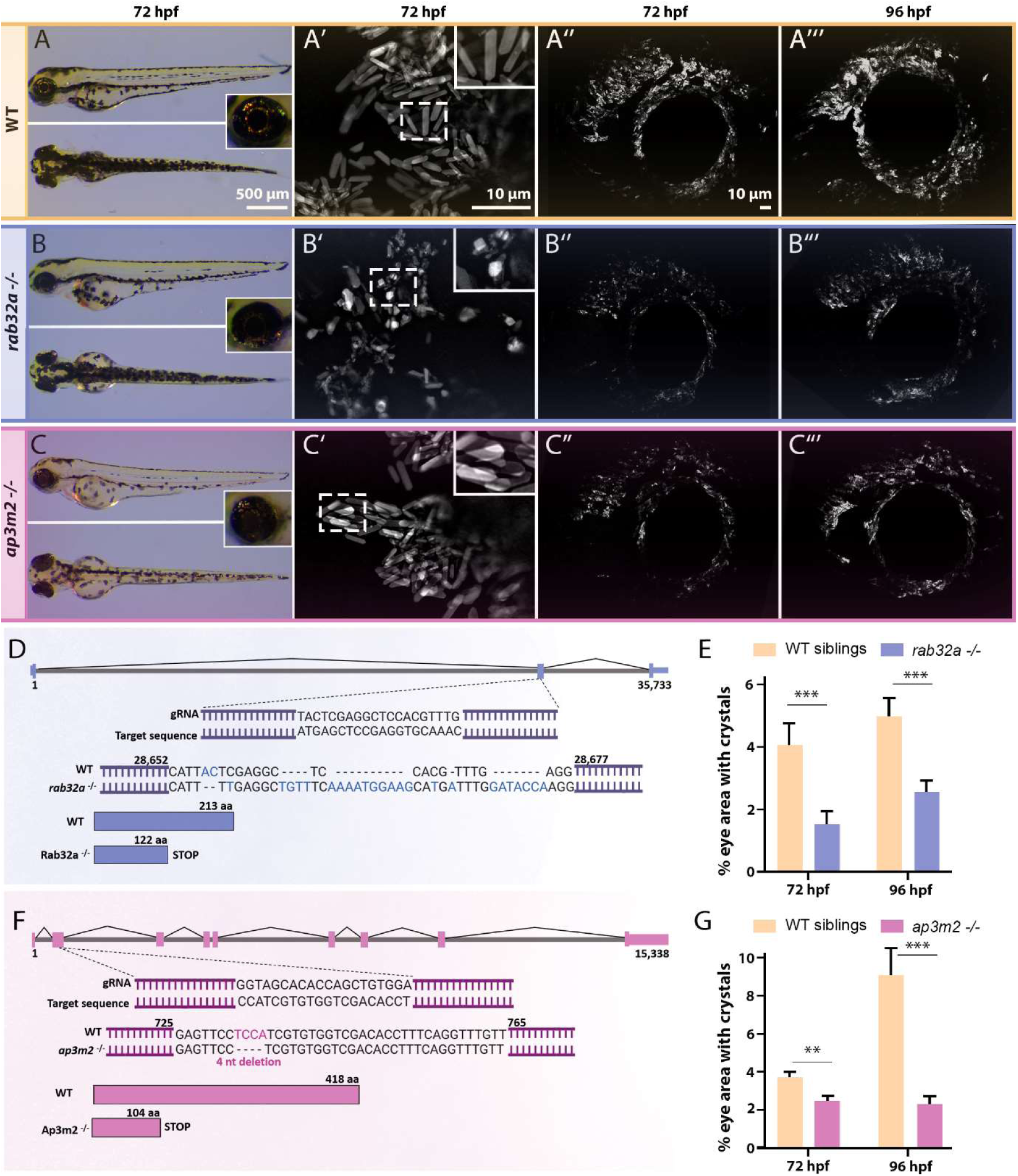
Reduced crystal production in *rab32a* and *ap3m2* mutants, with distinct morphological alterations in *rab32a* mutants. (**A–C**) Brightfield images comparing WT (A) *rab32a^−/−^* (B), and *ap3m2^−/−^* (C) larvae at 72 hpf. Insets highlight decreased iris reflectivity in *rab32a^−/−^* and *ap3m2^−/−^* mutants relative to WT. *ap3m2*^−/−^ mutants also display reduced melanophore pigmentation. (**A′–C′**) Confocal MIPs of iridophores from WT and mutants at 72 and 96 hpf. Both mutants exhibit reduced crystal accumulation compared to WT. Insets in A′ and B′ highlight shorter and distorted crystals in *rab32a^−/−^* mutants, while *ap3m2^−/−^* mutants maintain WT-like crystal morphology. (**D, F**) Schematic of CRISPR-Cas9 targeting strategy for *rab32a* and *ap3m2*. *rab32a^−/−^* mutants harbor frameshift-inducing mutations leading to premature stop codons; *ap3m2^−/−^* mutants carry a 4-nt deletion resulting in truncation. Diagrams partially generated using BioRender.com. (**E, G**) Quantification of the percentage of eye area occupied by crystals in WT siblings versus mutants at 72 and 96 hpf. Data represent mean ± s.e.m. from ≥3 independent clutches. (E) 72 hpf: WT, n = 17; *rab32a^−/−^*, n = 18. 96 hpf: WT, n = 18; *rab32a^−/−^*, n = 21. (G) 72 hpf: WT, n = 21; *ap3m2^−/−^*, n = 31. 96 hpf: WT, n = 18; *ap3m2^−/−^*, n = 17. Significance determined by two-way ANOVA: ***P < 0.001; **P < 0.01; *P < 0.05. n=number of total larvae measured in different biological repeats.

Both homozygous *rab32a*^−/−^ and *ap3m2*^−/−^ mutants showed a marked reduction in the number of crystals formed by iridophores (**Fig. 3**), without any detectable effect on overall larval development (**Supplementary Fig. 3A, B**). A similar phenotype was observed in knockdown experiments using morpholinos (**Supplementary Fig. 2**), thereby further validating their roles in iridosome development. The specific expression of *rab32a* and *ap3m2* genes in iridophores was also verified using a hybridization chain reaction (HCR) assay (**Supplementary Fig. 3A-B**).

Interestingly, while the crystals formed by *ap3m2*^−/−^ mutants were similar in morphology to those found in wild-type (WT) fish (**Fig. 3A’ and C’**), *rab32a*^−/−^ mutants produced significantly shorter, with lower aspect ratio and more distorted crystals (**Fig. 3B’ and Supplementary Fig. 3I**). These findings indicate that both RAB32a and AP3M2 are crucial for iridosome biogenesis and crystal production. RAB32a appears to play a particularly significant role in proper crystal shaping and structural integrity, whereas AP3M2 is more specifically involved in organelle biogenesis and crystal quantity.

### Severe disruption of iridosome biogenesis and crystal formation in *hps5* mutants

To further assess the role of LRO machinery in iridosome biogenesis, we investigated *hps5*, a component of the BLOC-2 complex identified in the third node of our network analysis (**Fig. 4A–D**). In melanophores, BLOC-2 mediates trafficking of cargo such as TYRP1 from the trans-Golgi network to stage III melanosomes ^22,68,69^. Strikingly, *hps5*^−/−^ mutants exhibited a severely deficient capacity to form guanine crystals, with almost no crystals present in the eyes of 72 hours post fertilization (hpf) larvae (**Fig. 4 B and B’**) without any detectable effect on overall larval development (**Supplementary Fig. 3C**). High magnification of crystals in 7 dpf mutant larvae revealed irregular-shaped assemblies (**Fig. 4B’’**). The reduction in crystal formation and crystal shape impairment in the *hps5^−/−^* mutant was far more pronounced compared to the other mutants we analyzed, suggesting that iridosome biogenesis and maturation are severely impaired in *hps5^−/−^* mutants.

**Figure 4.**
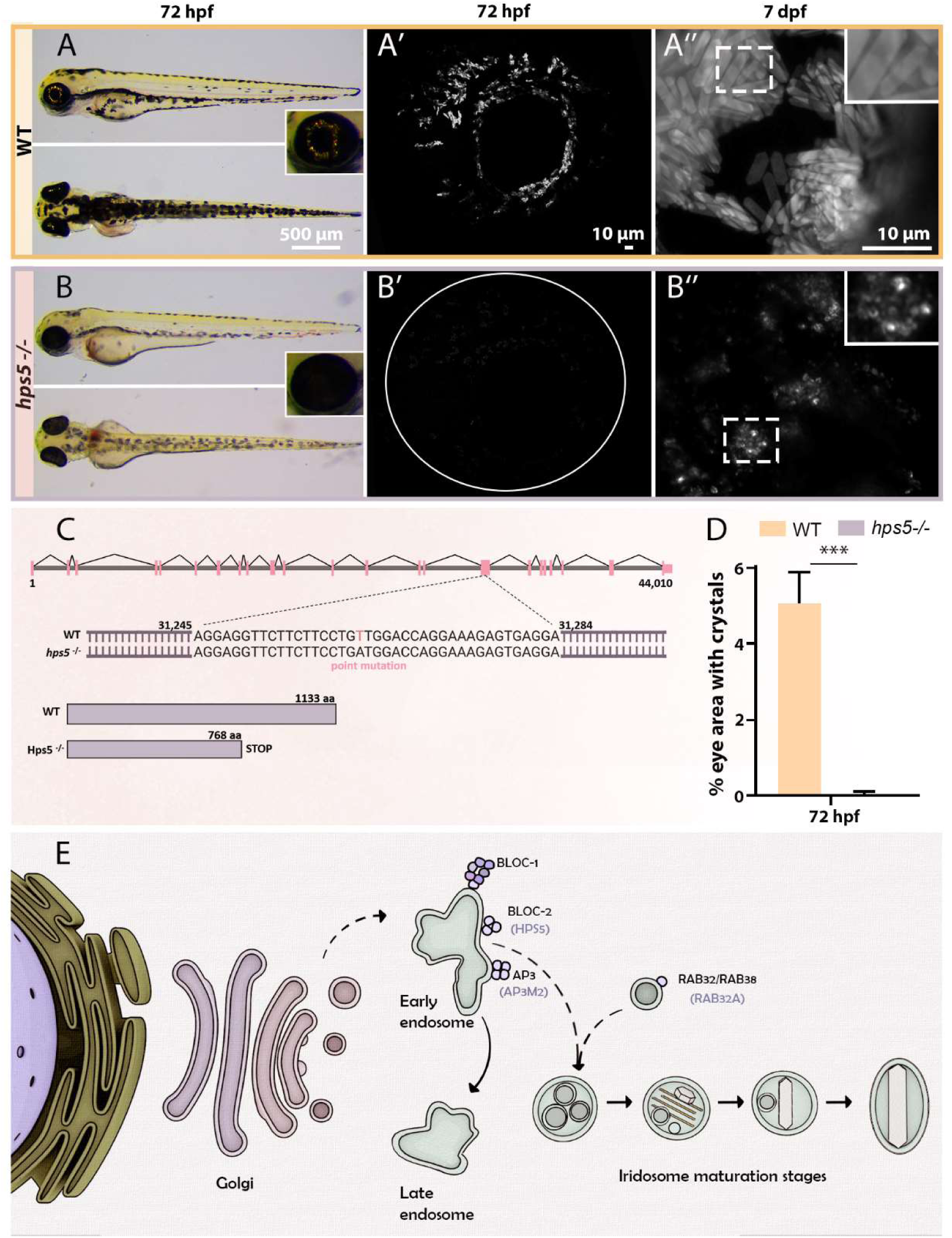
Severe disruption of iridosome biogenesis and crystal formation in *hps5* mutants. (**A–B**) Brightfield images of WT (A) and *hps5^−/−^* (B) zebrafish larvae at 72 hpf. Insets show lack of reflective crystals in *hps5^−/−^* eyes and reduced melanophore pigmentation compared to WT. (**A′–B′**) Confocal MIPs of iridophores at 72 hpf and 7 dpf showing near-complete absence of crystal formation in *hps5^−/−^* mutants. Insets highlight aberrant crystal aggregates in mutant iridophores. (**C**) Schematic of the *hps5* mutation, which introduces a premature stop codon resulting in a truncated protein. Illustration created in part using BioRender.com. (**D**) Quantification of the percentage of eye area containing crystals in WT and *hps5^−/−^* larvae at 72 hpf. Data represent mean ± s.e.m. from two clutches (WT, n = 10; *hps5^−/−^*, n = 9). Significance assessed by two-way ANOVA: ***P < 0.001. n=number of total larvae measured in different biological repeats. **(E)** Proposed model illustrating the involvement of the endolysosomal pathway and the trans-Golgi network in iridosome biogenesis. LRO components implicated in crystal formation - including RAB32a, AP3M2, and HPS5 - are indicated in brackets.

### The presence of intraluminal vesicles within iridosomes suggests an endosomal origin

After establishing the identity of iridosomes as LROs, we next sought to investigate their structural maturation and cellular origin using cryogenic electron tomogrpahy. Early-stage iridosomes contained multiple thin, developing crystals, whereas more mature iridosomes typically housed a single, larger crystal (**Supplementary Fig. 4**), consistent with previous observations. Notably, we frequently observed the presence of intraluminal vesicles (ILVs) adjacent to the growing guanine crystals in maturing iridosomes (**Supplementary Fig. 4**).

The detection of ILVs within iridosomes provides important mechanistic insights. ILVs are a hallmark of multivesicular bodies (MVBs) and are characteristically found during the maturation of multiple LRO types, including melanosomes and cytotoxic granules^21,24,25^. Their presence within iridosomes strongly suggests that these organelles originate from, or transit through, an early or late endosomal intermediate during their biogenesis and maturation. Thus, iridosomes not only share molecular signatures with classical LROs but also display structural hallmarks indicative of an endosomal origin.

### Iridosomes are conserved as LROs across diverse vertebrate lineages

To investigate whether the LRO identity of iridosomes is conserved beyond zebrafish, we analysed publicly available single cell RNAseq datasets from other vertebrate species. Specifically, we calculated an “LRO score” for every iridophore cells from two additional organisms: the medaka (*Oryzias latipes*)^16^, a teleost related to zebrafish, and the leopard gecko (*Eublepharis macularius*)^46^, a distantly related reptile (**Supplementary Fig. 5 and Supplementary Table. 1**). Using the same 27-gene LRO signature applied to zebrafish, we found that both medaka and gecko iridophores exhibited significantly elevated LRO scores compared to their respective general cell populations (Medaka: Student’s t-test, two tailed, unpaired, P< 0.0001; Gecko: Two-way ANOVA followed by Tukey analysis. P< 0.0001 for iridophores versus non pigment cells and melanophores versus non pigment cells. P< 0.05 for iridophores versus melanophores).These results further support the notion that the molecular machinery underlying iridosome biogenesis is deeply conserved across vertebrate taxa (**Supplementary Fig. 5**).

## Discussion

### Iridosomes as a new class of lysosome-related organelle and their molecular pathways

Our findings establish that iridosomes, the guanine crystal-bearing organelles of zebrafish iridophores, are a previously unrecognized member of the LRO family. By integrating transcriptomic profiling, gene perturbation, cryo-TEM, and comparative evolutionary analyses, we reveal that iridosome biogenesis depends on a similar molecular machinery that governs the formation of canonical LROs such as melanosomes, platelet dense granules, and Weibel-Palade bodies^29,30^ (**Fig. 4E**). These results place iridosomes within the expanding paradigm of LRO diversity and highlight their potential as a model for studying specialized organelle biogenesis.

Iridosomes align with LROs in several key respects. First, we show that iridophores express a suite of LRO-associated genes, including *rab32*, BLOC complex components, adaptor protein complexes such as AP3, and vacuolar sorting and proton pump machinery. This constellation of molecular players is a hallmark of the LRO class. Second, our functional studies demonstrate that disrupting key LRO regulators-RAB32a, AP3M2, and HPS5, impairs crystal formation and/or organelle maturation. These defects recapitulate the phenotypes observed in canonical LRO systems, such as melanosome or dense granule disorders in Hermansky-Pudlak syndrome.

Notably, the phenotypes of *rab32a* and *hps5* mutants suggest distinct contributions to crystal morphology and organelle maturation, mirroring the specialized trafficking roles previously ascribed to these proteins in melanosomes^64,68–71^. In melanosomes, RAB32 facilitate the delivery of cargo to maturing organelles through interactions with BLOC complexes^39,64,70^. Their loss leads to mislocalization of cargo proteins and impaired pigment synthesis^64,70^. Similarly, our findings suggest that RAB32a is required in iridophores to direct the trafficking of specific components necessary for proper crystal morphogenesis, such as guanine transporters, scaffold proteins, or pH-regulating channels, thus influencing crystal shape and organization. HPS5 functions in the BLOC2 complex, which in melanophores is involved in transport from early endosomes to maturating melanosomes^22,38,39,69,71^. Strikingly, *hps5* mutants had a severely deficient capacity to form guanine crystals, and almost no crystals were observed in the eyes of larvae. This reduction in crystal formation was much more pronounced than any of the other mutants we previously studied, suggesting that iridosme biogenesis is severely defective in these mutants. Together, these results point to a functional division of labor among LRO regulators during iridosome development, in which certain components control organelle identity and cargo input, while others modulate the physical characteristics of the organelle’s contents. This mechanistic architecture closely parallels with what has been described in melanosome biogenesis and further supports the classification of iridosomes as a specialized subtype of LROs.

Beyond the core LRO machinery shared with melanosomes and other systems, our transcriptomic analysis also identified several candidate regulators - *rab20*, *rab3il1*, and *rabl6b,* that may be unique to iridophores. These genes are not known to participate in melanosome biogenesis, suggesting that they may confer additional trafficking or regulatory capabilities tailored to the demands of guanine crystal formation. For instance, they could facilitate the transport of guanine precursors, modulate the luminal environment, or coordinate interactions between iridosomes and other organelles. Their selective enrichment highlights a possible layer of organelle-specific specialization within the broader LRO framework and raises the possibility that iridosomes have evolved dedicated molecular tools to support their crystalline function.

### Endosomal origin, pH dynamics and intraluminal vesicle architecture

Beyond their molecular signature, iridosomes display defining structural features of endosomal origin. Cryo-TEM imaging revealed the presence of intraluminal vesicles (ILVs) within maturing iridosomes - a hallmark of multivesicular intermediates observed in other LROs^24,25^. In melanosomes, for example, ILVs serve as organizing centers for PMEL amyloid fibrils that template melanin deposition.^72^ By analogy, the ILVs observed in iridosomes may provide organizing centers for similar fibers or may serve for nucleation centres for guanine crystallization by themselves.

Another unifying feature of LROs is the modulation of their luminal pH, which regulates cargo condensation, enzymatic activity, and scaffold maturation^29,34,40^. In melanosomes, lumen acidification is essential for tyrosinase activation and early pigment polymerization^41,73^. Our previous work demonstrating early iridosome^42^ acidification, together with our current transcriptomic evidence for upregulated vacuolar-type H⁺-ATPases, suggests that iridosomes follow a similar developmental trajectory. We propose that nascent iridosomes are acidified to concentrate guanine precursors or control their solubility, with subsequent pH neutralization enabling crystal nucleation and elongation. The observation of internal, sheet-like or fibrillar structures in iridosomes from zebrafish and invertebrates (e.g., scallops, spiders)^26,28,42,74^ further supports a model in which LRO-derived scaffolds and acid–base transitions coordinate precise biomineralization within this specialized organelle.

### Evolutionary conservation and expansion of the functional spectrum of LROs

The evolutionary conservation of iridosome identity as an LRO is further supported by our cross-species meta-analysis. We show that both the medaka and zebrafish, that are separated by approximately 200 million years of evolution, as well as reptiles, which diverged from teleost fish around 400 million years ago^75^, express orthologs of key LRO regulators in their iridophores. This suggests that iridosomes have ancient evolutionary roots, with their formation relying on a conserved regulatory network. This widespread conservation implies strong selective pressure to maintain crystalline optical organelles for light manipulation, camouflage, or signaling across environments and taxa.

Our results also broaden the functional scope of LROs to include crystal forming organelles, adding a new dimension to the known functional diversity within this organelle class. While most LROs store pigments, small molecules, or secreted factors, iridosomes leverage LRO machinery to orchestrate spatially precise crystallization of purine metabolites. This finding reinforces the idea that LROs are not defined by their contents, but rather by their shared reliance on a specialized endolysosomal trafficking and maturation system tailored to cell-type-specific roles.

### Iridophores as a model for organelle morphogenesis and bioinspired design

Iridophores and their iridosomes offer an exceptional model for studying rapid organelle biogenesis, membrane-based crystal nucleation, and lumenal self-assembly processes. Given the tractability of the zebrafish model and the optical readouts provided by iridophores, future studies could leverage iridosomes to dissect how cells coordinate trafficking, pH control, and scaffold deposition to drive organelle specialization.

Finally, the ability of iridophores to achieve precise control over guanine crystal morphology and reflectivity may hold translational promise for materials science. As bioinspired optics and photonic devices increasingly look to nature for design principles, understanding the cell biological basis of iridosome biogenesis could guide the engineering of tunable, light-manipulating nanostructures.

In summary, this study classifies iridosomes as lysosome-related organelles and reveals a conserved molecular framework for their biogenesis. These findings not only advance our understanding of pigment cell diversity and endomembrane specialization but also open new avenues for investigating biomineralization and cellular optics through the lens of organelle biology.

## Materials and Methods

### Zebrafish Husbandry and Handling

Zebrafish (*Danio rerio*) were housed at ∼28 °C, 14 hour light:10 hour dark and fed with Artemia and flake food. Fish were maintained and fed following the standard protocols. Crosses were performed with at least 3 old adults. Embryos were kept in E3 zebrafish embryo medium at 28 °C until reaching the desired developmental stage. Lines that were used in this work are as follows: Et(Ola.Edar:GAL4,14xUAS:GFP)^TDL358Et^ (RRID: ZDB-FISH-150901-2380)^77^, denominated ET358 for simplicity, Tg(*pnp4a*:PALM-mCherry)^wprt10Tg^ (RRID: ZDB-FISH-210414-18)^51^, *hps5^-/-^* mutant (sa37981, European Zebrafish Resource Center (EZRC)) (RRID: ZDB-ALT-160601-5792) and AB (WT).

### Cell Dissociation

Cell dissociation was carried out by customizing a protocol described previously^26,27,42^. Briefly, Tg(*pnp4a*:PALM-mCherry) positive 5dpf larvae were anesthetized with Tricaine (0.1% MS-222 (Sigma)) and immersed in TrypLE Express (Invitrogen, 12604039). Fish were incubated at 37°C and shaken at 200 rpm for 2 hr, followed by mechanical disruption with vortex and pestels to further dissociate the cells. Cells were then strained through a 40 µM cell strainer with cold HL-15 buffer with 1mM EDTA and centrifuged at 3000 RCF for 5 min at 4°C (HL-15 Buffer: Hank’s Balanced salt solution 40% (Sigma H8264) and 60% Leibovitz’s L-15 Medium (Gibco 21083-027)). The pelleted cells were then resuspended in 10 ml of fresh cold HL-15 with 1mM EDTA and then were incubated with Hoechst (H3570, Invitrogen)) to mark the nuclei for 30 min before FACS.

### FACs Sorting

Cell isolation was carried out by customizing a protocol described previously^26,27,42^. Cells were isolated from Tg(*pnp4a*:PALM-mCherry) positive fish larvae and sorted via Fluorescence-Activated Cell Sorting (FACS). Cells were analyzed and sorted using a BD FACSAria™ III or V Cell Sorter with a 100 µM nozzle. Cells were illuminated using a both 405 and 561 nm lasers. Cells were gated based on attributes to separate cells from each other as well as from cellular debris. Cellular debris were detected using forward, side scatter and Hoescht signals to select against the smallest particles (1 mm or less). Cells were additionally sorted and enriched based on detection using 561 nm filters, corresponding to the *pnp4a*:PALM-mCherry signal. Cells (∼20,000 cells per sample) were collected directly into lysis buffer (Dynabeads® mRNA DIRECT™ Purification Kit (Ambion, 61012)). After reaching ∼20,000 cells each sample was vortexed, spinned down and placed immediately on dry ice. The cell collection was performed in three biological replicates and stored in −80°C before proceeding to next steps of sample preparation. For cryo-TEM cells were collected into ice-cold HL-15 medium with 1% FBS and kept on ice until mounted on grids for downstream imaging.

### RNA isolation

Purification of RNA from sorted cells (∼20,000 iridophores or general cell population) was done using Dynabeads® mRNA DIRECT™ Purification Kit (Ambion, 61012) according to manufacturer protocol. The purified RNA samples were frozen at −80°C until the step of RNA-seq library preparation.

### Bulk MARS-Seq preparation

RNA-seq libraries were prepared at the Crown Genomics Institute of the Nancy and Stephen Grand Israel National Center for Personalized Medicine, Weizmann Institute of Science. A bulk adaptation of the MARS-Seq protocol^78,79^ was used to generate RNA-Seq libraries for expression profiling of iridophores and general control cell population. Briefly, RNA (all RNA amount purified from ∼20,000 sorted cells) from each sample was barcoded during reverse transcription and pooled. Following Agencourct Ampure XP beads cleanup (Beckman Coulter), the pooled samples underwent second-strand synthesis and were linearly amplified by T7 in vitro transcription. The resulting RNA was fragmented and converted into a sequencing-ready library by tagging the samples with Illumina sequences during ligation, RT, and PCR. Libraries were quantified by Qubit and TapeStation as well as by qPCR as previously described^78,79^. For *Danio rerio,* the housekeeping gene *actinb1* was selected for its relatively high and uniform expression^53^ and was specifically calibrated for the MARS-Seq protocol. Sequencing was done on a NextSeq 500 using NextSeq High Output, 75 cycles, allocating 400M reads in total (Illumina).

### Bioinformatics pipeline methods

#### RNAseq analysis

Data was analyzed with the UTAP pipeline (v1.10.2)^80^. The pipeline steps included fastq trimming using Cutadapt (DOI: 10.14806/ej.17.1.200) with the following parameters: -a AGATCGGAAGAGCACACGTCTGAACTCCAGTCAC -a “A” --times 2 -u 3 -u −3 -q 20 -m 25. The trimmed reads were then mapped to the danRer11 genome using STAR (v2.5.2b). Quantification of raw read counts per gene was performed using the UCSC gene annotation. The quantification process involved marking read duplicates with an in-house UTAP script, followed by quantification using HTSeq-count (DOI: 10.1093/bioinformatics/btu638) in union mode. As MARseq is a 3’ protocol, the quantification was applied to a region that includes the 3’ 1000 bp of each transcript and 100 bases downstream of the 3’ end was used for quantification.

Read count normalization and differential expression analysis were performed using DESeq2 (v1.36.1) (DOI: 10.1186/s13059-014-0550-8), with the betaPrior parameter set to TRUE. Genes with a log2 fold change (log2FC) > 1, an adjusted p-value (p.adj) < 0.05, and an avregare of > 10 normalized read count in at least one of the compared conditions were considered differentially expressed.

The genes si:ch1073-75o15.4 and LOC100301575 are scaled outside of the plot therefore they were not included in the volcano plot.

#### scRNAseq analysis

##### Leopard Gecko

- The raw gene count matrix from the study^46^ (GSE264342) was downloaded from GEO and reanalyzed using Seurat v5.1.0, and the same parameters and procedures as described in ^46^. Doublet detection was performed using scDblFinder (v1.18.0), and cells identified as doublets were excluded from further analysis. Low-quality cells were filtered out based on the following criteria: fewer than 1,000 detected genes, fewer than 2,000 UMIs, more than 9,000 detected genes, or more than 70,000 UMIs. To eliminate stressed or dying cells, we also removed cells with over 10% mitochondrial gene expression and over 30% ribosomal gene expression (63.3% of the cells passed these filters). Normalization was performed using Seurat’s log-normalization method. Scaling was applied while regressing out cell cycle effects, using human cell cycle gene sets provided by Seurat.

Following dimensionality reduction by PCA, we performed unsupervised clustering using Seurat’s default Louvain algorithm at a resolution of 0.8 on a shared nearest neighbor graph constructed from the first 20 principal components. Nonlinear dimensional reduction for visualization was performed with the RunUMAP function using the first 20 principal components. Marker genes were identified using the FindAllMarkers function with the Wilcoxon rank-sum test, applying min.pct = 0.2 and logfc.threshold = 0.2. Genes with adjusted p-values < 0.05 (Bonferroni correction) were considered significant. Annotation of the different cell populations was based on the expression of the following genes derived from Figure S15 of : COL1A1, TWIST1, DSP, LOC129338255, EDAR, SOX2, TEK, PECAM1, MYOD1, TYRP1, OCA2, DCT, TYR, CORIN, SOX10, MITF, PMEL, LOC129342328, PAX7, LTK, TFEC, PNP, ALX4, and LOC129340104. Focusing on pigment cells, we used PNP expression as a marker for iridophores, OCA2 as a marker for melanophores. The annotation was further validated by querying the PanglaoDB database with the marker genes of each cluster, using the R package, enrichR^81^.

LRO score of each cell was calculated using the AddModuleScore function of Seurat, and with the genes listed in the supplementary table 1. These are genes with a well-established function in Lysosome-related organelles.

##### Medaka

- Medaka scRNAseq data were available from the study of ^16^. LRO score was calculated using the AddModuleScore function of Seurat, and with the genes listed in Supplementary Table 1.

#### Metascape analysis

Gene pathways enrichment analysis for identifying upregulated biological processes for identified genes was done using a Metascape algorithm^58^ (http://metascape.org). As a background for the analysis we used all genes that were detected in the data.

#### *rab32a* and *ap3m2* CRISPR mutants

The *rab32a* and *ap3m2* CRISPR mutants were generated as described previously^82^. Briefly, trans-activating crispr RNAs (crRNAs) were designed for specific loci of the *rab32a* and *ap3m2* genes using the predesigned crRNAs dataset from Integrated DNA Technologies (IDT). Several crRNAs were tested for ribonucleoprotein (RNP) mutagenesis and were chosen considering where the start codon is located, as well as high on-target and low off-target scores. The most efficient crRNA tested was located on exon 2 of the *rab32a* and exon 2 of the *ap3m2* canonical transcripts, with target sequence 5′-TACTCGAGGCTCCACGTTTG-3′ (Dr.Cas9.Rab32A.1.AA, IDT) 5′-CCATCGTGTGGTCGACACCT −3′ and (Dr.Cas9.AP3M2.1.AD, IDT) respectively. Each tested crRNA was separately annealed with an equal molar amount of tracrRNA (1072533, IDT) and diluted to 57 μM in Duplex buffer (11-01-03-01, IDT), generating the single guide RNA (sgRNA). The RNP mixes were assembled using Cas9 protein (Alt-R S.p. Cas9 Nuclease V3; 1081058, IDT; 61 μM stock) and a sgRNA, in equimolar amounts, generating a 28.5 μM RNP solution. One-cell-stage WT AB embryos were injected with 1 nl of each of the RNP mixes. To screen for the efficiency of RNP mutagenesis pools of 5 embryos of each RNP-injected condition were individually genotyped. Founder fish (F0) containing frameshift mutations were identified by genotyping the resulting F1 progeny, resulting from outcrosses with WT AB fish. F1 progeny were genotyped and fish carrying identical mutations crossed to produce F2 generation that contained mutant, WT siblings and heterozygous fish. F3 pure mutant line was raised after crossing F2 mutant fish.

The mutant lines that were generated: (1) *rab32a*^-/-^ resulting in a series of genetic alternations (see Fig. 3), leading to a truncated protein, resulting in 122 aa instead of 213 aa. (2) *ap3m2*^-/-^resulting in a 4 nt deletion (see Fig.3), leading to a truncated protein, resulting in 104 aa instead of 418 aa.

#### Genotyping

Genomic DNA from individual larvae, pool of larvae or adult after fin clipping, was extracted according to following protocol: samples were incubated in 100 µl 50 mM NaOH at 95°C for 20 minutes then cooled in 4°C for 2 minutes. 10 µl of Tris-HCL (pH 8) was added to each sample that was vortexed and centrifuged at 12,000 rcf for 2 minutes. This was followed by performing PCR with Red Load Taq Master mix (Larova) with primers surrounding the *rab32a* mutation region (forward, 5′ -TTATGTCCCACGTCTAGGTCAG-3′; reverse, 5′ -GCTTCGGCTTGACCTATTAC −3′) or with primers surrounding the *ap3m2* mutation region (forward, 5′ -TGTAAGGATGCTGGCAGGTTG-3′; reverse, 5′ - CCTGGCTACAGGCCTAATTA −3′). For genotyping of *hps5*^-/-^ (sa37981, European Zebrafish Resource Center [EZRC]) adult fish and larvae the following primers were used surrounding the *hps5* mutation region in exon 16 (forward, 5′ - TAGAGAGAGCACCCCCTGTTCCATC-3′; reverse, 5′ - TACAGTAAACTCACCCAGCATGCCA −3′). The point mutation results in nonsense mutation leading to a truncated protein, resulting in 768 aa instead of 1133 aa (see Fig.4). All PCR products were sent for sanger sequencing. Sequences were analyzed using Snapgene software.

#### Live Imaging

For imaging, live larvae were anesthetized with Tricaine and then mounted with 0.9 % low-melting-point agarose (Sigma, A9414) on an imaging glass bottom dish (Cellvis, d35-14-1.5-n). Live imaging was carried out on an inverted Zeiss LSM 980 or 900 confocal microscopes using X20, X40 or X63 lenses. Crystals were imaged with reflection. For time-lapse imaging, mounted larvae were covered with E3 medium with tricaine and placed inside a chamber with controlled 28°C. Imaging was done up to 48 hours. Images from each session were processed into a movie using the ImageJ software.

For confocal imaging quantifications, Z-stack images of the eye area with crystals (1.25 µm each slice, total 30-50 µm) were taken using X20 magnification lens. Whole larvae imaging for standard length (SL)^83^ measurements were done using X2.5 lens.

For gross morphology images, larvae were anesthetized with Tricaine and placed in Methyl Cellulose and imaged with Leica M80.

For high resolution images of the crystals (Fig. 3 A’, B’, C’ and Fig. 4 A’’, B’’), live larvae were mounted on glass slides and their eyes imaged with X100 lens using the Leica DM-6B upright microscope. The images were then analysed using ImageJ for length and width measurements of individual crystals and the aspect ratio calculated for each crystal.

#### Plunging and Cryo-TEM

As previously described^26,27^, sorted iridophore cells (3.5 µl) with 15 nm gold beads (1 µl) were applied to glow-discharged holey carbon R2/2 Cu 200 SiO_2_ mesh grids (Quantifoil) coated with collagen, Type I, rat tail (EMD Millipore, 08-115) for cell adherence. The grids were blotted and vitrified by plunging into liquid ethane using a Leica EM GP automatic plunger at 4 °C and 90% humidity. Frozen grids were kept in liquid nitrogen until used. Data were collected on a Titan Krios TEM G3i (Thermo Fisher Scientific) equipped with a BioQuantum energy filter and a K3 direct electron detector (Gatan). Cryogenic electron tomography data (shown in **Supplementary** Figure 4) was collected at 300 kV with the K3 camera (counting mode) using Thermo Fisher Tomography software. The TEM magnification corresponded to a camera pixel size of 1.6 Å, and the target defocus was set to −3 μm. The total dose for a full tilt series was 140 electrons per Å^2^. Tomogram tilt series was collected using the dose-symmetric scheme, ±60° at 2° degrees steps. The tilt series images alignment and reconstruction were performed in IMOD^84^.

#### Hybridization chain reaction fluorescence in situ hybridization (HCR) assay

5dpf WT (AB) larvae were fixed in 4% PFA over night at 4°C, washed 3 times in PBT (Triton 0.1% in PBS) and transferred to 100% MeOH to be stored in −20°C until further use. For HCR assay, fixed larvae were washed twice in SSCTX2 and ∼10 larvae per sample were incubated with 4nM probe (*pnp4a* and *rab32a* or *ap3m2*, Molecular Instruments) in preheated hybridization buffer (2XSSCT,10% dextran sulfate, 10% formamide, Molecular Instruments) at 37°C, overnight with agitation. Next samples were washed with preheated wash buffer (2XSSCT, 30% formamide, Molecular Instruments) for 30 min at 37°C with agitation, then washed twice for 20 min in SSCTX5 at RT and incubated for 30 min in amplification buffer (5XSSCT,10% dextran sulfate, Molecular Instruments) at RT. Next, samples were incubated with appropriate snap-cooled hairpins (hairpins h1and h2 (Molecular Instruments) 180nM each) in amplification buffer overnight in the dark at RT, followed by washes with SSCTX5 (3X 20 min) at RT. The samples were then mounted and imaged using confocal microscopy for RNA localization in the larval eye.

#### Microinjection of embryos with morpholinos

One-cell-stage TDL358 or *pnp4a*:PALM-mCherry embryos were injected using standard procedures with a PV850 microinjector and M3301 Manual Micromanipulator (World Precision Instruments) that were used for microinjection. Gene knockdown experiments were performed using the following morpholino-modified antisense oligonucleotides (MO, Gene Tools):

*rab32a*, 5′ TGCCATGCTGCCAAAATGTCAGTTA 3′; AUG MO.

*ap3m*2, 5′ ACTAACAGAGATTACCTGTGATGGT 3′; Exon Junction MO. control morpholino, 5′ CCTCTTACCTCAGTTACAATTTATA 3′ (Gene Tools stock control).

Each oligonucleotide was diluted with ultrapure (Milli-Q) water to 1 mM and 1–2 pmol of MO was injected into one-cell-stage embryos. Embryos were left to develop at 28.5 °C until the desired stage and then live-imaged.

#### Reflectance and Fluorescence Coverage Quantification

For the quantification step done on confocal images, a custom graphical user interface application was developed. The application accepts images in the czi format and creates maximal intensity projections over the images, allowing for further processing and visualization. The application then performs an automatic object detection and segmentation step, followed by optional manual annotation, user specified thresholding and area to area calculation and output. In every step of the workflow the application allows visualization of the dataset and of the segmentation. It was implemented in Python, utilizing OpenCV, PIL, Tkinter/CustomTkinter, Matplotlib, NumPy, and Pandas for image processing, visualization, and data management, and with aicspylibczi for handling czi files. The application was specifically designed for the lab and is available on Gur-Lab-WIS GitHub page.

#### Statistical Analysis

Statistical experimental details can be found in the text and in the relevant figure legends. Differences were considered significant if the P value was < 0.05 (*), < 0.01 (**), < 0.001 (***) or < 0.0001 (****). Non-significant results were marked as > 0.05 (ns). Two-way ANOVA, testing for genotype and batch and unpaired two-tailed Student’s t-tests were performed where appropriate to compare group means. In **Supplementary Fig. 5**, data from medaka was subjected to Student’s t-test, two tailed, unpaired and the data from gecko was analysed with two-way ANOVA followed by Tukey analysis. Tests were run using R, v. 4.4.2. Aspect ratios were compared between genotypes using a linear mixed effects model, with genotype as a fixed factor and fish ID as a random factor. Analyses were done using the ‘lmerTest’ package in R, v. 4.5.0

## Supporting information

RNA seq data

## Data, Materials, and Software Availability

Output of bioinformatic analysis is available as Datasets S1. RNA-seq GEO Accession number: <u>GSE297759</u> (Enter token inifsssafpmpvgx into the box). All other data are included in the article and/or supporting information.

## Acknowledgments

We thank Yuval Elazari (Crown Genomics Institute, Nancy and Stephen Grand Israel National Center for Personalized Medicine, Weizmann Institute of Science) for assistance with RNA-seq data processing. We are grateful to Ziv Porat (Life Sciences Core Facilities, Weizmann Institute of Science) for his guidance in developing and optimizing the FACS sorting protocol. We thank Neta Varsano (Department of Chemical Research Support, Weizmann Institute of Science) for her contributions to illustration and figure layout design. We also thank Ron Rotkopf (Life Sciences Core Facilities, Weizmann Institute of Science) for his support with statistical analysis and data interpretation. This work was supported by an ERC Starting Grant (Grant number: 101077470, “CRYSTALCELL”) and by the Israel Science Foundation (grant No. 691/22) awarded to D.G. Electron microscopy studies were conducted at the Irving and Cherna Moskowitz Center for Nano and Bio-Nano Imaging at the Weizmann Institute of Science. R.D. is a fellow of the Ariane de Rothschild Women Doctoral Program.

