## Supplementary Information for "Lysosome-Related Organelles Orchestrate Guanine Crystal Formation in Pigment Cells"

### Table of Contents:

Supplementary Figure 1. The iridophore transcriptome is enriched for lysosome-related organelle (LRO) components.

Supplementary Figure 2. Downregulation of *rab32a* and *ap3m2* impairs crystal formation in zebrafish iridophores.

Supplementary Figure 3. Mutations in *rab32a*, *ap3m2* or *hps5* genes have no effect on SL.

Supplementary Figure 4. Intraluminal vesicles are present in developing iridosomes.

Supplementary Figure 5. Iridosomes are conserved as LROs among vertebrates.

Supplementary Table 1. LRO marker set genes list.

Supplementary Movie 1.

Supplementary Movie 2.

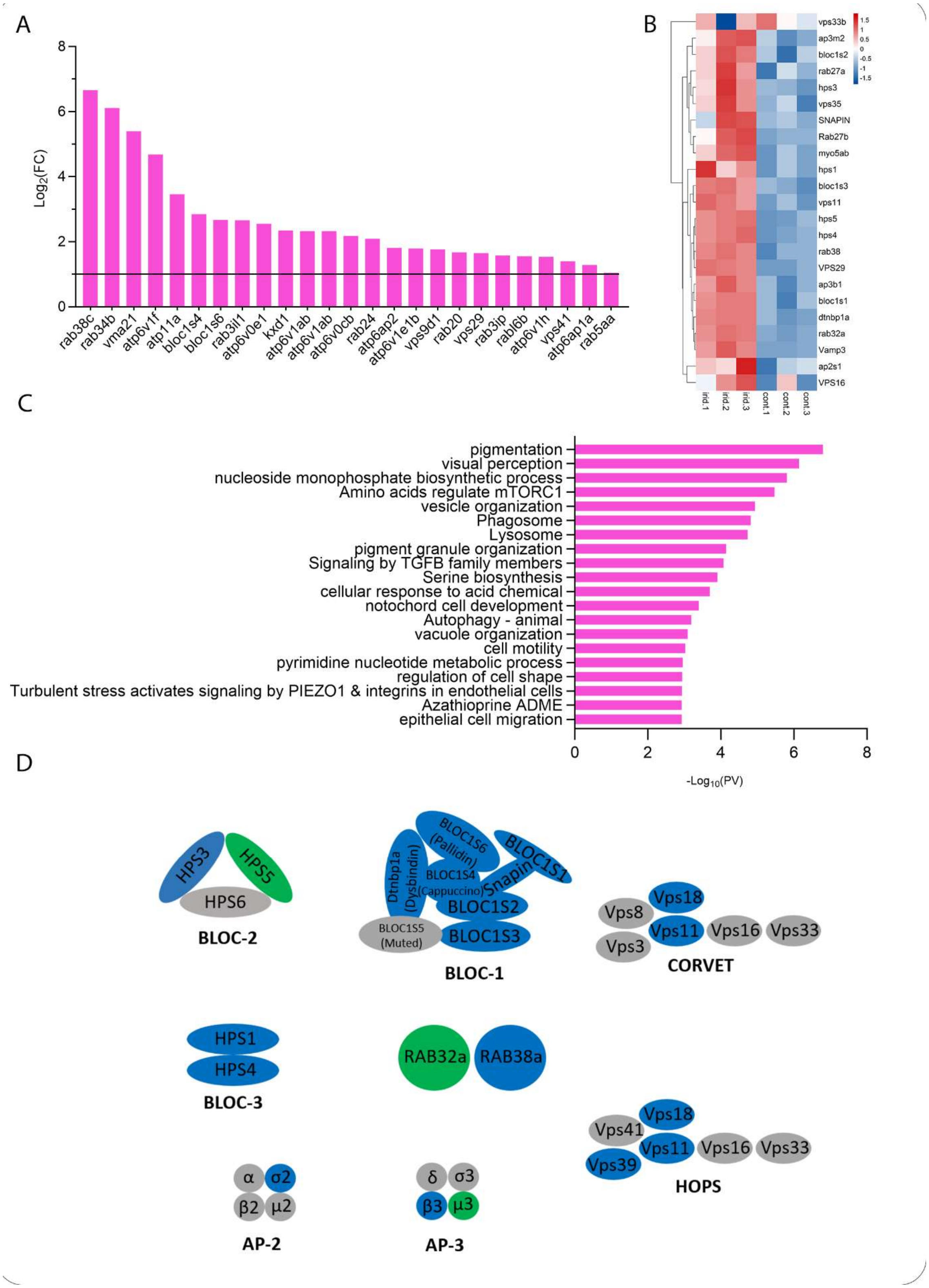


**Supplementary Figure 1. The iridophore transcriptome is enriched for lysosome-related organelle (LRO) components.** (**A**) Bar graph showing elevated expression levels of additional genes involved in LRO biogenesis and V-ATPase subunits significantly upregulated in iridophores. (**B**) Heatmap depicting the expression of the LRO marker set genes across biological replicates of sorted iridophores (irid) and control cell populations (cont) from the Bulk MARS-Seq experiment. The expression in the heatmap is scaled per every gene, the plot was made with the package pheatmap of R. (**C**) Full results of functional enrichment analysis performed with Metascape[58], highlighting biological processes significantly enriched among upregulated genes (pink). The entire list of genes expressed was used as a background in the analysis. (**D**) Diagrams of key LRO-associated proteins and complexes. Genes upregulated in iridophores are highlighted in blue, and those functionally validated in this study are marked in green. Mammalian gene names written in brackets.

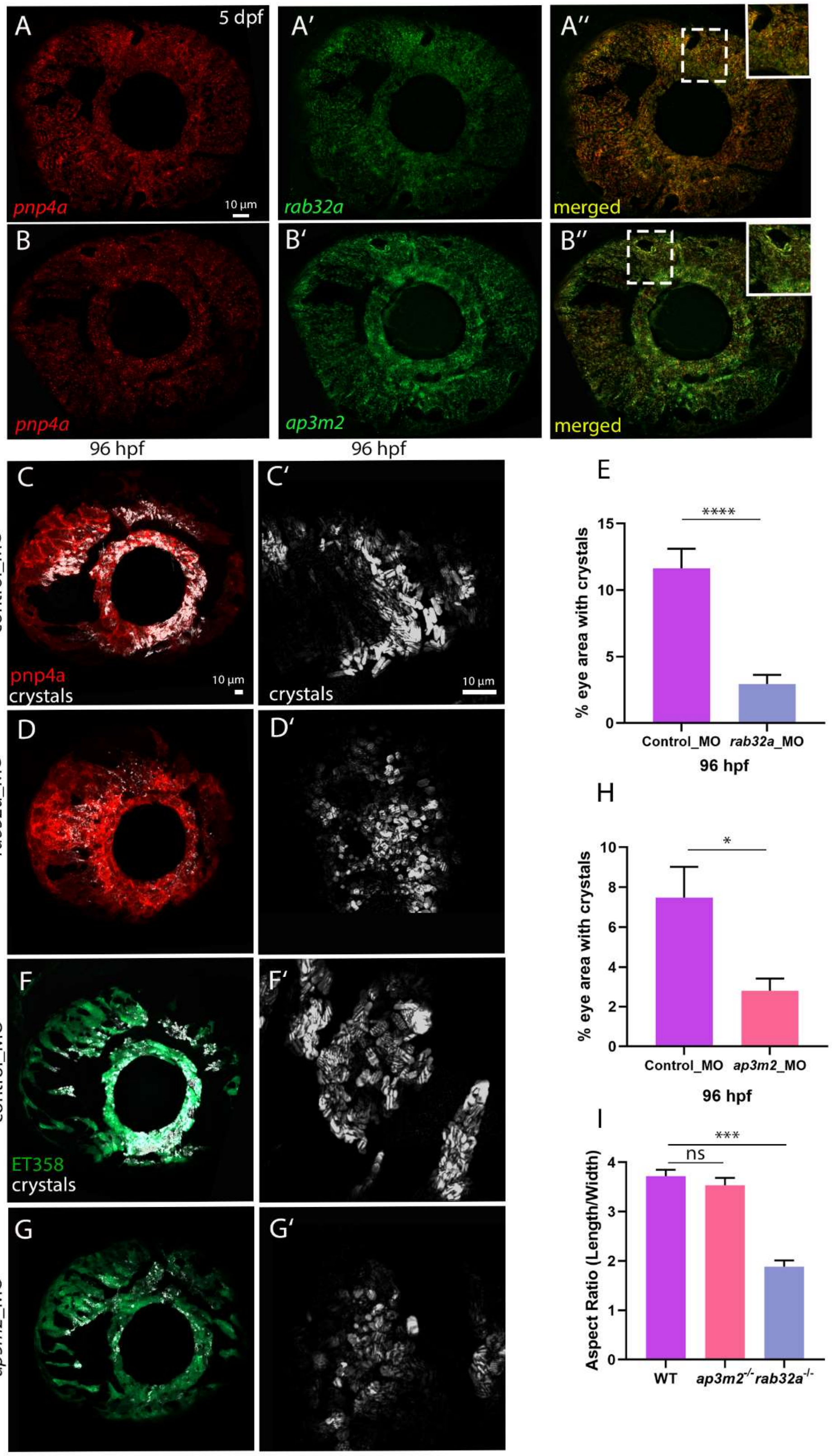
A
5 dpf
pnp4a
10 µm
A′
rab32a
A″
merged
B
pnp4a
B′
ap3m2
B″
merged
96 hpf
96 hpf
C
control_MO
pnp4a
crystals
10 µm
C′
crystals
10 µm
D
rab32a_MO
D′
F
control_MO
ET358
crystals
F′
G
ap3m2_MO
G′
E
****
% eye area with crystals
15
10
5
0
Control_MO
rab32a_MO
96 hpf
H
*
% eye area with crystals
10
8
6
4
2
0
Control_MO
ap3m2_MO
96 hpf
I
***
ns
Aspect Ratio (Length/Width)
4
3
2
1
0
WT
ap3m2-/-
rab32a-/-

**Supplementary Figure 2. Downregulation of *rab32a* and *ap3m2* impairs crystal formation in zebrafish iridophores.** (**A–B**) Confocal maximum intensity projections (MIPs) of 5 dpf larval eyes following hybridization chain reaction (HCR) assay. mRNA localization of *rab32a* (A) and *ap3m2* (B) is shown in green; iridophores are marked by *pnp4a* transcript in red. Co-localization of transcripts (A″, B″ and insets) confirms *rab32a* and *ap3m2* expression in iridophores at this developmental stage. (**C-D, F-H**) Confocal images showing results of morpholino (MO)-mediated knockdown of *rab32a* and *ap3m2*. Iridophores are labeled with either *pnp4a*:PALM-mCherry (red) or ET358:GFP (green), and guanine crystal reflectance is shown in white. (**C–E**) Knockdown of *rab32a* results in decreased crystal quantity and abnormal morphology (D′) compared to the control. (**E**) Quantification of the percentage of eye area covered by crystals in control_MO and *rab32a*_MO larvae at 96 hpf. Data represent mean ± s.e.m. from two independent experiments (control_MO, n = 9; *rab32a*_MO, n = 13). Statistical significance determined by two-way ANOVA; ****P < 0.0001. (**F–H**) Knockdown of *ap3m2* leads to a moderate reduction in crystal number with minimal impact on morphology compared to the control. (H) Quantification of the percentage of eye area occupied by crystals in control_MO and *ap3m2*_MO larvae at 96 hpf. Data represent mean ± s.e.m. from one biological replicate (control_MO, n = 6; *ap3m2*_MO, n = 7). Statistical analysis performed using two-tailed unpaired Student's t-test; *P < 0.05. n=number of total larvae measured in different biological repeats. **(I)** Aspect ratio (length/width) of individual crystals from eyes of WT, *ap3m2*$^{-/-}$ and *rab32a*$^{-/-}$ 72 hpf larvae. Data represent mean ± s.e.m. from three biological repeats (WT, n = 60; *ap3m2*$^{-/-}$ , n= 40; *rab32a*$^{-/-}$, n =45).; Aspect ratios were compared between genotypes using a linear mixed effects model, with genotype as a fixed factor and larvae ID as a random factor. ***P < 0.001, ns= non significant (P>0.05). n=number of crystals measured in different biological repeats.

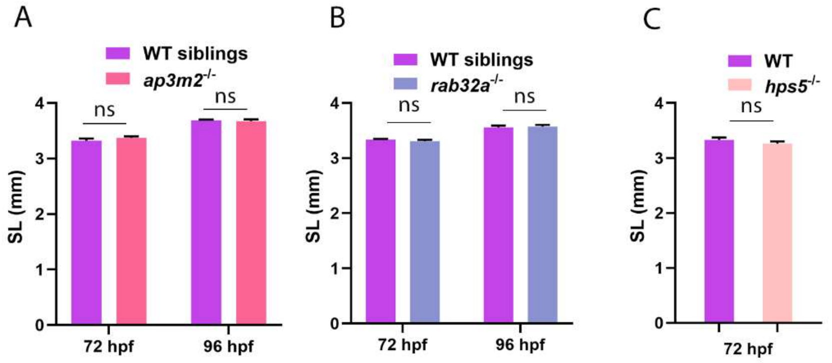


**Supplementary Figure 3. Mutations in *rab32a*, *ap3m2* or *hps5* genes have no effect on SL. (A-C)** Quantification of standard length SL[83] (mm) in larvae of WT siblings and *rab32a*$^{-/-}$, *ap3m2*$^{-/-}$ and *hps5*$^{-/-}$ mutants during development (72 hpf and 96 hpf). The mean ± s.e.m. are shown for at least two independent biological repeats; (A) 72hpf: WT, n =17 ; *rab32a*$^{-/-}$ n=18 , 96 hpf: WT, n =18 ; *rab32a*$^{-/-}$, n=21. (B) 72 hpf: WT, n =21 ; *ap3m2*$^{-/-}$ , n=31 , 96 hpf: WT, n =18 ; *ap3m2*$^{-/-}$ n=17. (C) WT, n =10; *hps5*$^{-/-}$, n=9. Two-way ANOVA; ns= non significant (P>0.05). n=number of total larvae measured in different biological repeats.

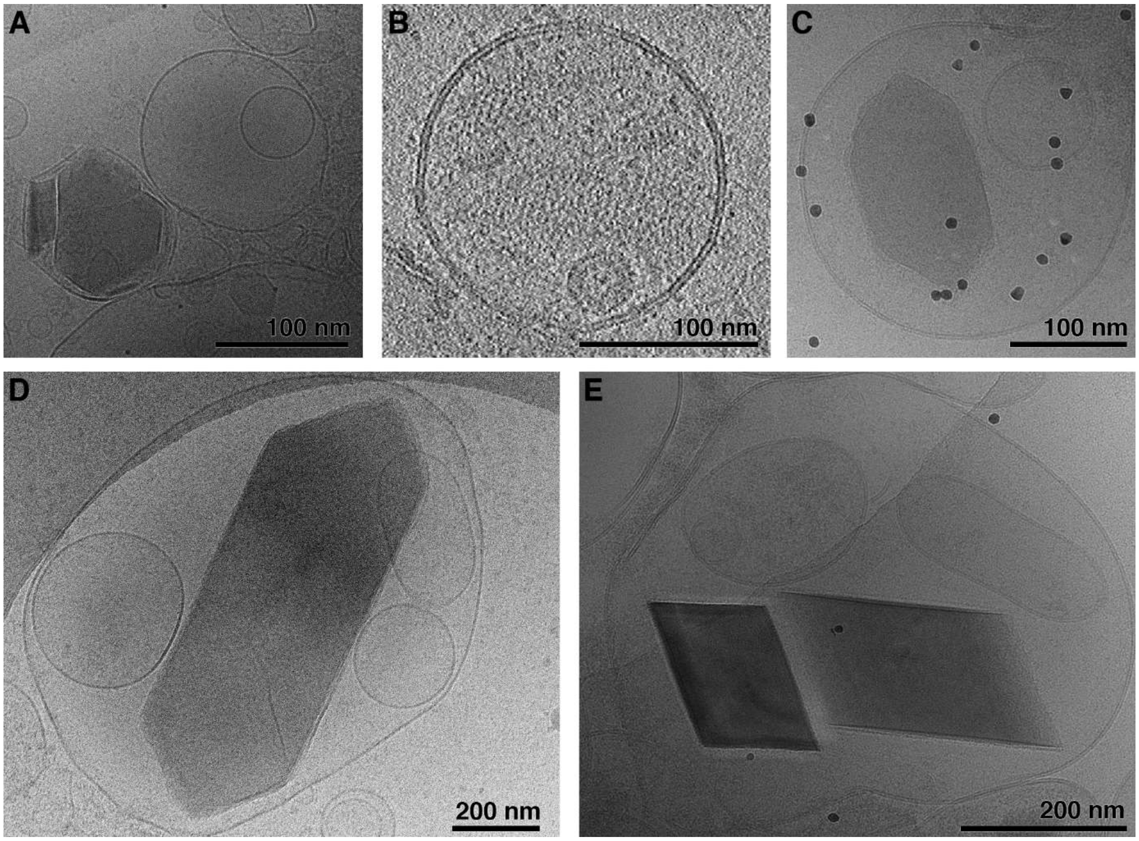


**Supplementary Figure 4. Intraluminal vesicles are present in developing iridosomes.** (**A–E**) Cryogenic electron microscopy images showing iridosomes at various stages of maturation within iridophores isolated from 72 hpf zebrafish larvae. (A) Early-stage iridosomes. (B) Cryogenic electron tomography reconstruction of Iridosome containing protein fibers and ILV. (C–E) Iridosomes at different maturation levels showing ILVs. High-contrast puncta visible in panels B and D correspond to 15 nm gold fiducials.

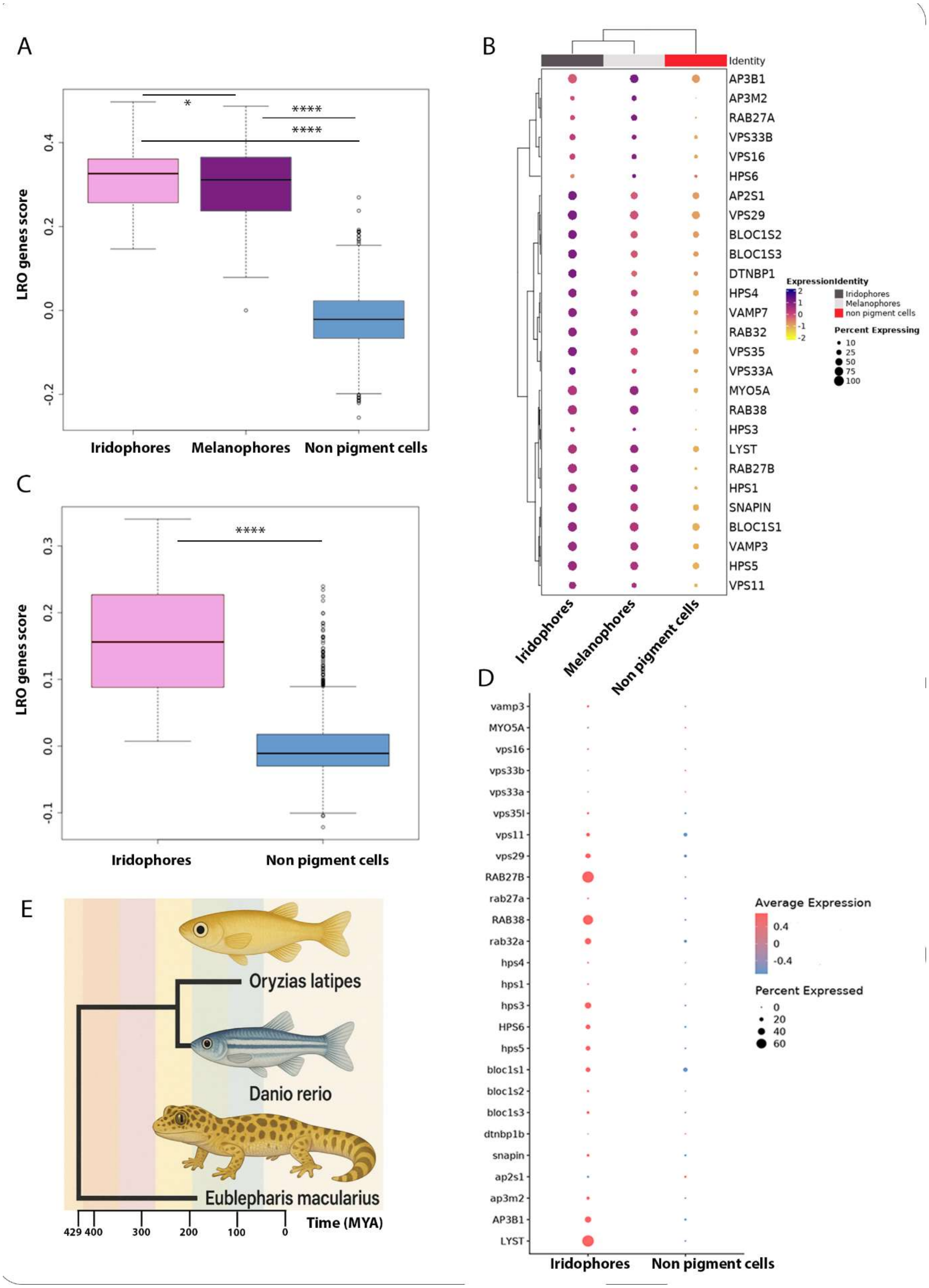


**Supplementary Figure 5. Iridosomes are conserved as LROs across vertebrates.**
(**A, C**) Bar plots showing the calculated "LRO score" for iridophore and melanophore populations compared to non-pigmented cells in the leopard gecko (*Eublepharis macularius*) (A) and medaka (*Oryzias latipes*) (C). LRO scores were derived from the average expression of a consensus set of LRO-associated genes. (**B, D**) Dot plots displaying the expression levels

of individual LRO marker genes in iridophores and non-pigmented cells from the leopard gecko (B) and medaka (D), demonstrating enriched expression in pigment cell types. Statistical analysis: medaka, unpaired two-tailed Student's t-test; ****P < 0.0001. Gecko: two-way ANOVA followed by Tukey post hoc test; ****P < 0.0001 for iridophores vs. non-pigmented cells and melanophores vs. non-pigmented cells; *P < 0.05 for iridophores vs. melanophores. (**E**) Phylogenetic tree depicting[75] the evolutionary relationships between zebrafish (*Danio rerio*), medaka (*Oryzias latipes*), and leopard gecko (*Eublepharis macularius*), generated using https://timetree.org based on estimated divergence times and modified for improved visualization.

| Zebrafish symbol | Gecko symbol | Medaka Symbol |
|---|---|---|
| *lyst* | *lyst* | *lyst* |
| *ap3b1* | *ap3b1* | *ap3b1* |
| *ap3m2* | *ap3m2* | *ap3m2* |
| *ap2s1* | *ap2s1* | *ap2s1* |
| *snapin* | *snapin* | *snapin* |
| *dtnbp1a* | *dtnbp1* | *dtnbp1b* |
| *bloc1s3* | *bloc1s3* | *bloc1s3* |
| *bloc1s2* | *bloc1s2* | *bloc1s2* |
| *bloc1s1* | *bloc1s1* | *bloc1s1* |
| *hps5* | *hps5* | *hps5* |
| *hps6* | *hps6* | *hps6* |
| *hps3* | *hps3* | *hps3* |
| *hps1* | *hps1* | *hps1* |
| *hps4* | *hps4* | *hps4* |
| *rab32a* | *rab32* | *rab32a* |
| *rab38* | *rab38* | *rab38* |
| *rab27a* | *rab27a* | *rab27a* |
| *rab27b* | *rab27b* | *rab27b* |
| *vps29* | *vps29* | *vps29* |
| *vps11* | *vps11* | *vps11* |
| *vps35* | *vps35* | *vps35l* |
| *vps33a* | *vps33a* | *vps33a* |
| *vps33b* | *vps33b* | *vps33b* |
| *vps16* | *vps16* | *vps16* |
| *myo5ab* | *myo5a* | *myo5a* |
| *vamp3* | *vamp3* | *vamp3* |
| | *vamp7* | |

**Supplementary Table 1. LRO marker set genes list.** A list of consensus 27 LRO markers genes that were used to quantify the 'LRO score' for iridophores in zebrafish, medaka and gecko.

**Supplementary Movie 1. Time-lapse imaging of crystal formation dynamics in zebrafish eye iridophores.** A wild-type (AB strain) zebrafish larva at 52 hpf was imaged using confocal microscopy (40× objective, 2× digital zoom) to visualize guanine crystal formation within eye iridophores. Z-stacks comprising 25 optical sections (0.6 µm per step) were acquired every 5 minutes over a 17-hour period. MIPs were generated using ImageJ. Guanine crystals are visualized via reflectance and pseudocolored in blue**.**

**Supplementary Movie 2. Early eye development and iridosome formation in zebrafish larvae.** A Tg(*pnp4a*:PALM-mCherry) zebrafish larva at 51 hpf was imaged using confocal microscopy (20× objective, 1× digital zoom) to visualize iridophore development and crystal formation in the eye. A total of 66 Z-stacks (1.25 µm step size) were acquired every 26 minutes over a continuous 26-hour imaging session. MIPs were generated using ImageJ. Iridophore membranes are labeled in red (PALM-mCherry), and guanine crystals are visualized by reflectance and pseudocolored in blue.
